# Longitudinal monitoring of developmental plasticity in the mouse auditory cortex

**DOI:** 10.1101/2025.05.12.653595

**Authors:** Megan A. Kirchgessner, Mihir M. Vaze, Robert C. Froemke

## Abstract

The postnatal brain undergoes substantial plasticity in order to represent features and statistics of the sensory world. To date, for technical reasons it has been difficult to examine experience-dependent changes over the course of postnatal development. Here we perform longitudinal two-photon calcium imaging of hundreds of neurons in the auditory cortex of young mice, from postnatal day (P) 12 into adulthood. Auditory cortical neurons started responding to tonal stimuli by P13-14 with an initial tonotopic organization that expanded over the next few days. We documented the daily variation in tuning curves and best frequency, and while some neurons gradually changed their frequency representation across days, altogether our findings support the functional maturation of a largely stable auditory map at the population level. We then compared the representation of mouse pup ultrasonic vocalizations to pure tone responses. Ultrasonic pure tone tuning developed with a delay, coinciding with emerging responses to ultrasonic vocalizations. Vocalization responses initially were observed in neurons tuned to ultrasonic frequencies, but over later development the vocalization responses became independent from ultrasonic frequency tuning. Our results show how varied sensory representations at the single-cell and population levels in the postnatal auditory cortex emerge and change over the course of early-life development.

## Introduction

Sensory neurons represent and process information about the external world to govern perception and behavior. How that information is represented by individual neurons has been the topic of great interest and debate for over a century, since the first description of ‘receptive fields’ of isolated optic nerve fibers^1^. Sensory cortical neurons in early postnatal animals also have receptive fields, as described by Hubel and Wiesel in young, visually inexperienced kittens^2^. In the intervening years, studies in the visual cortex^3–7^, auditory cortex^8–10^, and somatosensory cortex^11^ found that neurons possess receptive fields soon after receiving sensory input, and specific sensory experience seems not to be required for the initial responses early in life. However, it has been difficult to determine how individual neuronal tuning profiles and receptive fields change during development and during early sensory experience, given the technical challenges of monitoring activity from individual identified neurons in young animals over prolonged timescales of days to weeks.

Pioneering developments in surgical, imaging, and data processing techniques have enabled the use of methods such as longitudinal two-photon imaging to record neuronal activity across early stages of postnatal development in rodents^12–16^. This approach has been used to monitor developmental changes in spontaneous and synchronous network activity patterns^11,12,17–22^ and to study sensory-evoked activity and emergence of receptive fields in the postnatal visual^5,7,23–26^, auditory^27,28^, and barrel cortices^11^. However, it remains unclear how the sensory responses of individual cortical neurons first emerge and are modified throughout life, especially as animals gain experience with behaviorally-relevant and more complex stimuli.

Vocalizations are one such class of ethologically relevant sensory cues that guide many aspects of behavior, such as courtship, parental care, and infant survival^29^. In rodents, vocalizations are spectrotemporally complex and generally emitted in the ultrasonic range^30,31^. In contrast to the visual cortex where receptive fields in neurons across the lifespan have been extensively characterized for both artificial and naturalistic stimulus classes^1,3,32,33^, satisfactory descriptions of auditory cortical neuron receptive fields that can explain neuronal responses across stimulus classes to include vocalizations have remained elusive^34^. Neuronal responses to vocalizations are not well predicted by their classical receptive fields for tonal stimuli^34^. For example, neurons in the mouse auditory cortex that respond to pup ultrasonic vocalizations often fail to respond to ultrasonic tonal stimuli^35–38,39^. Forms of experience-dependent plasticity (e.g., changes to lateral inhibition evoked by vocalizations) may account for the uncoupling between tuning to lower-level acoustic features and more complex auditory objects^37–40^. It remains unknown how neuronal selectivity for complex and important stimuli such as vocalizations emerges and changes over the course of life. This is required to understand how the cortical coding of complex stimuli relates to receptive field organization, and what aspects of sensory responses are learned or acquired by early experience versus being hard-wired or innate from birth.

## Results

### Two-photon calcium imaging of auditory cortical neuron activity over development

We injected C57Bl/6 neonatal mouse pups with a virus in the left auditory cortex to drive expression of the genetically encoded calcium indicator GCaMP8m^41^ in neurons under control of the CaMKII promoter (**Fig. 1a,b**). Juvenile pups underwent cranial window surgeries and headpost implants at P10-11 before daily imaging commenced at P12, corresponding to the approximate onset of hearing^42,43^. The same field of view of neurons (∼400-500 µm, average number of cells per field of view: 345±129 cells, mean±s.d.) was located each day based on the patterns of blood vessels and other cellular landmarks (**Fig. 1c-e**), and was imaged daily for at least the first week of hearing, with increasingly intermittent imaging thereafter for as long as animals remained healthy and the imaging window remained viable. There was a gradual decrease in the number of imaged cells across days (**Fig. 1e**; P12: 422±115 cells, P59-61: 272±206 cells, mean±s.d.), likely due to natural brain growth during this time period^14,44,45^. Despite the expansion of the skull and brain, we were able to image sound-evoked responses in the same field of view of neurons in the auditory cortex from the first week of hearing up until at least 6 weeks of age in 6/9 animals. We were also able to track individual neurons across development^14^ for 8-19 imaging sessions over 1-7 weeks (**Fig. 1f**).

**Figure 1.**
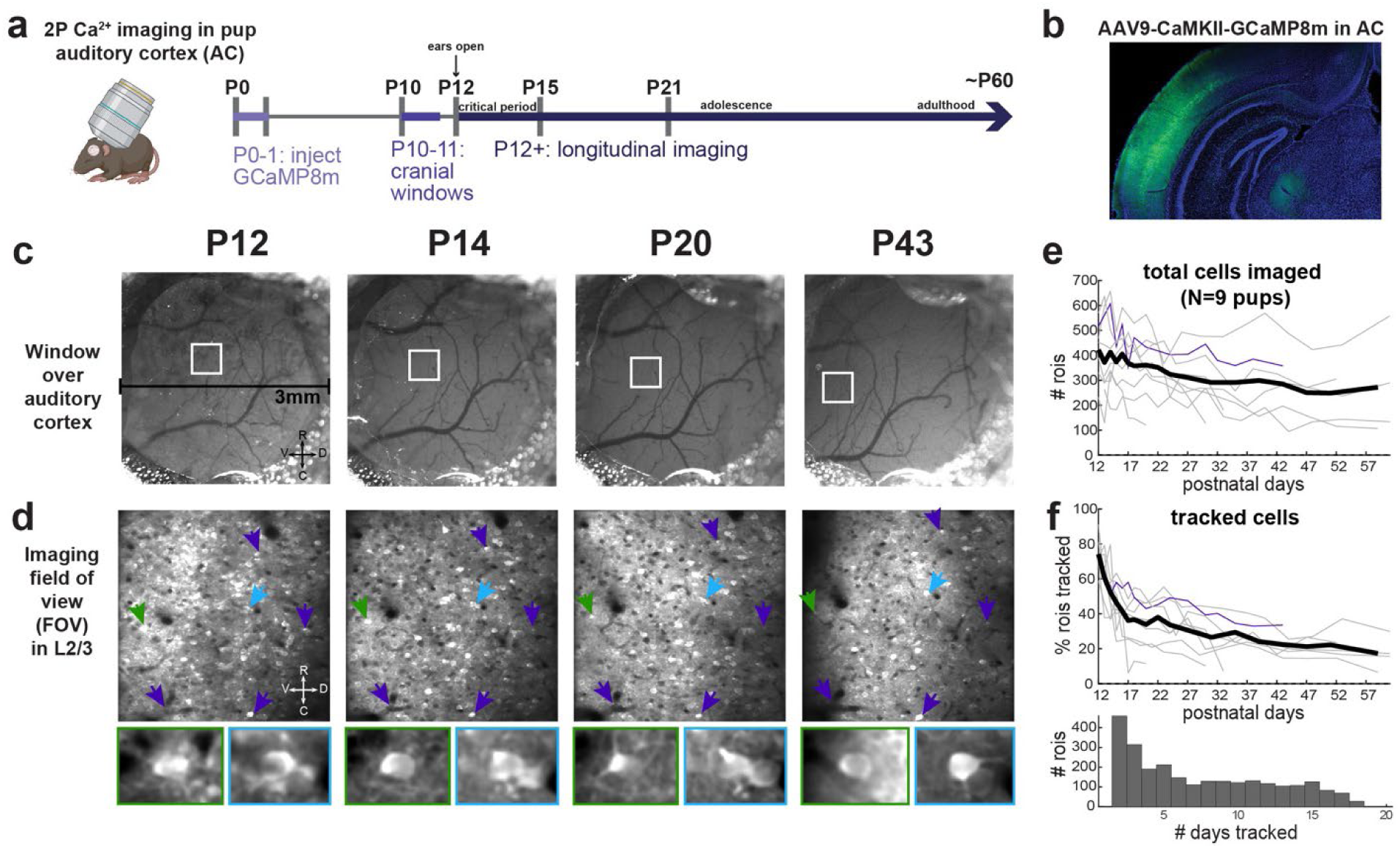
Longitudinal two-photon Ca^2+^ imaging to measure sound-evoked activity and tuning in developing mouse auditory cortex. **a**, Experimental timeline. **b**, GCaMP8m expression in the auditory cortex of a P60 animal, after neonatal AAV injection at P1 and cranial window insertion at P10. **c**, Widefield images of example window. **d**, Top, imaging field of view of layer 2/3 neurons. Arrows indicate cell landmarks across imaging days. Bottom, green and light blue-indicated cells across days. **e**, Number of ROIs (putative neurons) across imaging days for all animals (grey lines) and means across animals (black line). Purple line indicates example animal in **c,d**. **f**, Top, proportion of individual animal ROIs on each day that were tracked on at least one additional imaging day. Bottom, distribution of the number of tracked days for all tracked ROIs across animals (N=9 pups).

### Tonotopically organized sound-evoked activity emerges in the auditory cortex by P13-14

A substantial change in the activity patterns occurred between P12-14, with a reduction in spontaneous waves of activity transitioning to sound-evoked responses (**Fig. 2a**). On P12, reliable sound-evoked activation in L2/3 excitatory neurons of the left auditory cortex evoked by pure sine-wave tones (4-64 kHz, 70 dB SPL, half-octave spacing) or any other auditory stimuli was rare. Instead, there was a high degree of spontaneous, synchronous activations of spatially clustered groups of neurons, consistent with previous descriptions in other sensory cortical areas^11,17–19,46^.

**Figure 2.**
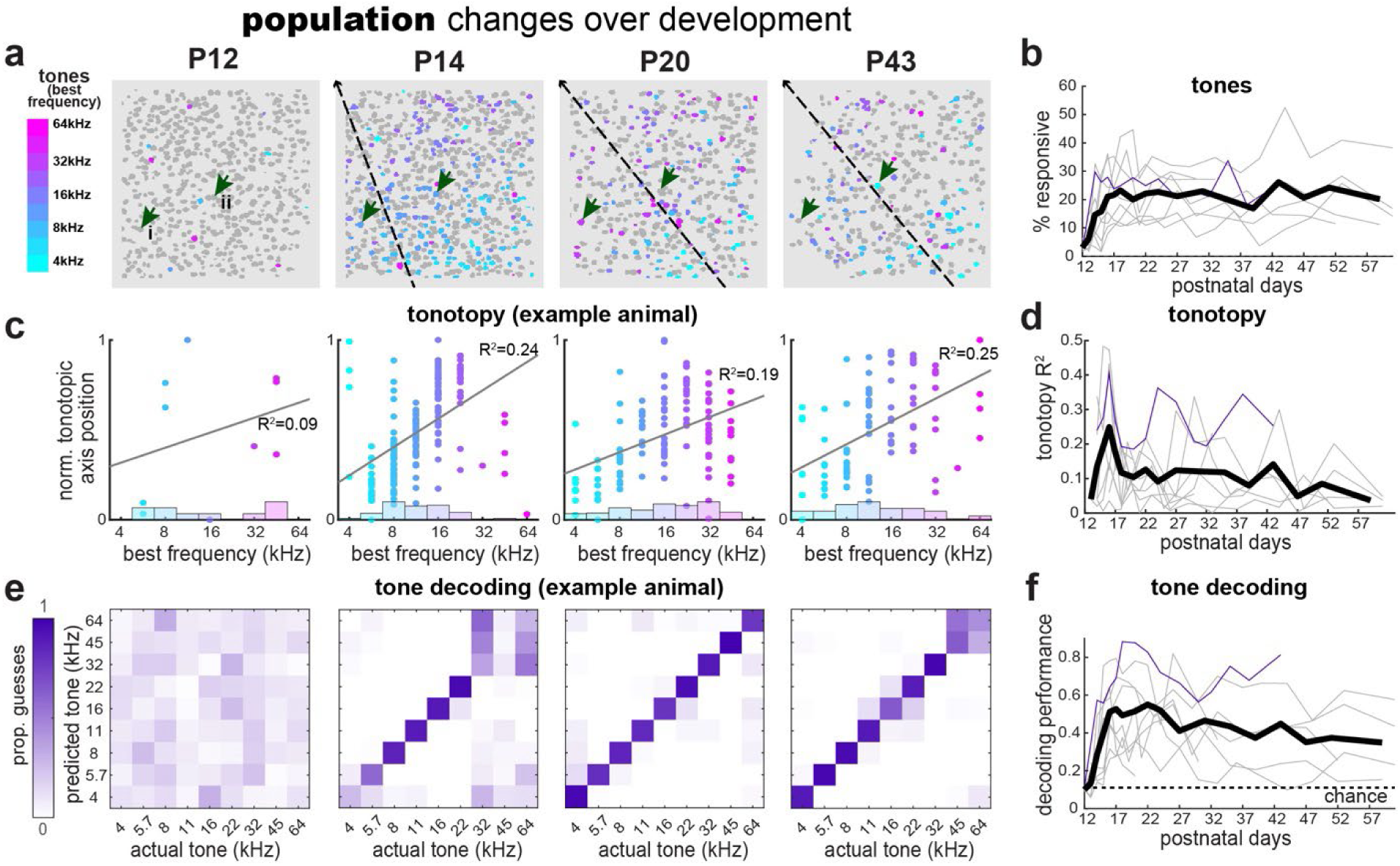
Population-level changes in tone-evoked activity in the developing auditory cortex. **a**, All Suite2p-extracted ROIs including significantly tone-responsive ROIs, colored according to the tone frequency eliciting the maximum median response (‘best frequency’, BF). Same example animal as in Fig. 1. Dashed lines indicate the inferred tonotopic axis. **b**, Proportion of significantly tone-responsive ROIs across postnatal days for all animals (grey = individual animals; purple = example animal in **a,c,e**; black = mean across N=9 animals). **c**, Correlation between each ROI’s BF and its position along the tonotopic axis. **d**, Tonotopic strength (R^2^ of correlation between BF and tonotopic position) across postnatal days. **e,f**, Linear classifier decoding of trial tone identity for example animal (**e**) and across days for all animals (**f**).

The proportion of tone-responsive cells in the left auditory cortex sharply increased at P13-14 (**Fig. 2a,b**). Tone-evoked activity in the postnatal auditory cortex was tonotopically organized, with individual best frequencies (i.e., tone eliciting largest median response) highly correlated with ROI position along the tonotopic axis (**Fig. 2c,d**). Tone-evoked responses were more consistent at P13 when higher intensity levels were used (85-95 dB SPL), likely due to initially high sensitivity thresholds of the cochlea during the first few days of hearing^43^. Over subsequent days of hearing, the range of best frequencies increased to encompass the full range of presented tones (4-64 kHz). The coding precision of cortical responses over days was quantified by the performance of a linear classifier trained to discriminate among the nine possible tone identities based on the population-level tone-evoked responses across trials (**Fig. 2e,f**). These results show that the mouse auditory cortex is tonotopically organized from the time of hearing onset and exhibits adult-like population activity and organization by P17-18, which is consistent with prior findings^8,10,47–49^.

### Stable versus shifting spectral tuning during the third postnatal week

Given the additional resolution afforded by longitudinal imaging and cell-tracking methodology^12,14^ for imaging the activity of individual neurons across days (**Fig. 1**), we sought to characterize the single-neuron changes that underlie these population-level changes in the functional maturation of the developing auditory cortex. Individual neurons could be tracked starting from the second postnatal week into adulthood (**Fig. 1f, 3a**). As we observed substantial population-level changes during the first week of hearing (**Fig. 2**), we focused our analysis on neurons that were tracked from the beginning (P13-15) to the end (P17-21) of this third postnatal week (PW3; n=1616 cells from N=9 juvenile pups). Some cells showed a gradual drift in frequency tuning towards higher frequencies in the third postnatal week (cell i, **Fig. 3a**), while others maintained consistent tuning across days (cell ii, **Fig. 3a**). During this time window, 41% of cells exhibited significant tone-evoked responses at any point: either only in the beginning of PW3 (‘lost response’, 11.0%), only at the end of PW3 (‘gained response’, 22.2%), or throughout PW3 (‘persistently responsive’, 7.9%; **Fig. 3b**). For ‘persistently responsive’ cells, we calculated the correlation between the tuning curves at the beginning versus the end of PW3. We observed varying degrees of shifting versus stable frequency tuning amongst individual cells, as indicated by the wide range in correlation coefficients (**Fig. 3c**). We also observed a range of increments and decrements of best frequencies, indicating significant heterogeneity in tuning changes (or lack thereof) across cells within the same field of views (**Fig. 3d**). However, it is possible that the direction of shifting tuning could depend on each cell’s initial tuning preference, which would result in an average change in best frequency close to zero as we observed. To examine this, we generated a transition probability matrix and found that on a day-to-day basis, cells were most likely to maintain the same best frequency, with some above-chance transitions towards slightly higher frequencies (**Fig. 3e**). Thus, while some single-cell tuning drift occurs to help drive the expansion of the tonotopic map, especially towards higher frequencies (**Fig. 2e,f**), tuning generally appeared to be quite stable, even during the first week of hearing.

**Figure 3.**
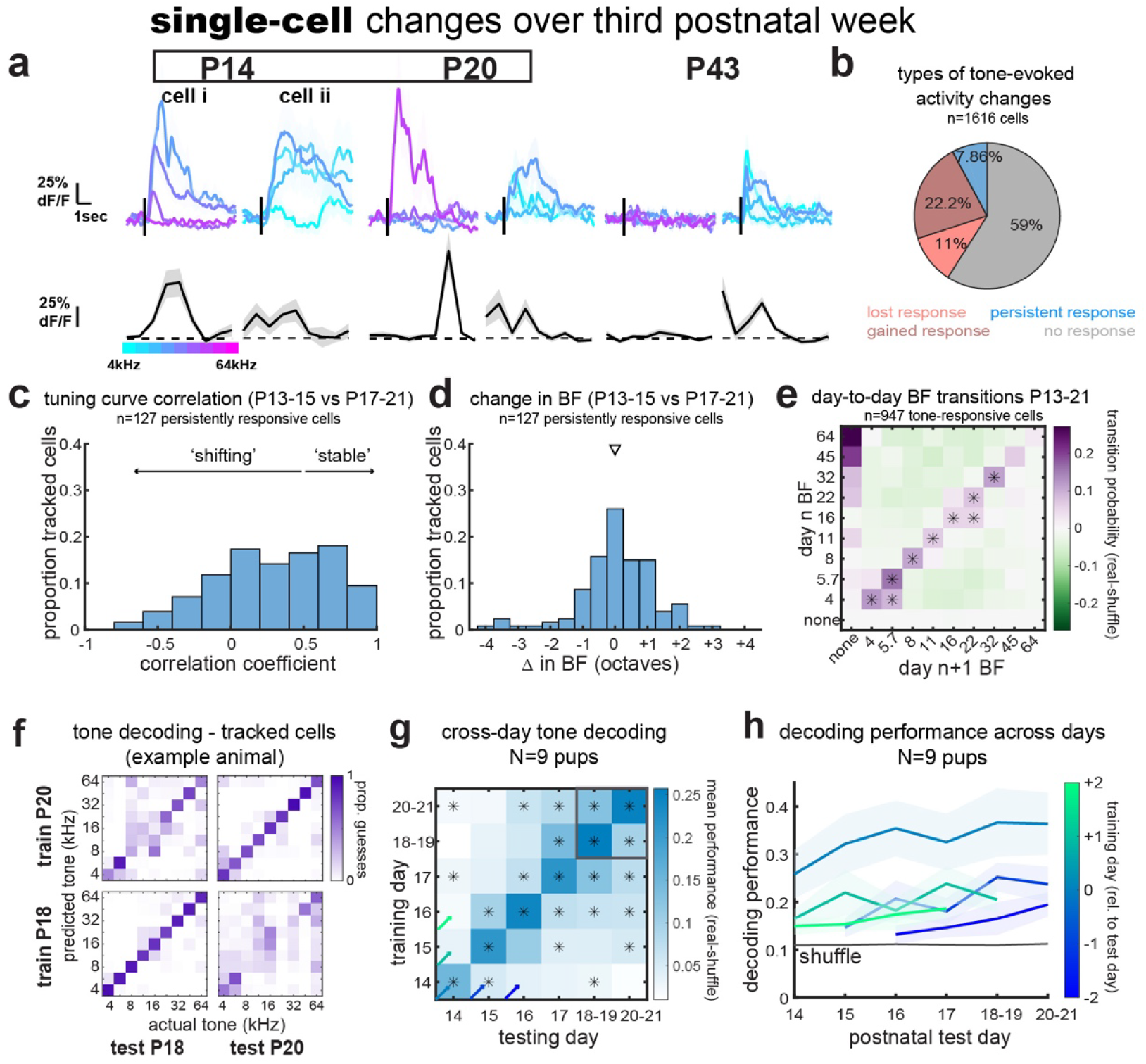
Single-cell changes in tone-evoked activity. **a**, Example cells tracked across days. Top, tone-evoked dF/F for example tones including best frequency (BF). Bottom, mean dF/F response by frequency. **b**, Proportion of response types among all cells tracked from P13-21. **c**, Correlation coefficients comparing tuning curves between P13-15 and P17-21. **d**, Change in best frequency from P13-15 to P17-21. Arrowhead, mean. **e**, Probability of day-to-day BF transitions for tone-responsive cells tracked for 2+ consecutive days. *p<0.05, permutation test with Benjamini-Hochburg FDR adjustment, q=0.025. **f**, Cross-day tone decoding from linear classifier trained and tested on tone-evoked activity of tracked neurons (same animal as **a**). **g**, Cross-day decoding performance (real-shuffled data) across animals. *p<0.05, q=0.025. **h**, Decoding performance (mean±s.e.m. across pups) over days for up to ±2 train-test day offsets (diagonals of matrix in **g**).

To better understand the stability of tone representations in the developing auditory cortex, we trained a linear classifier on the tone-evoked responses of our population of tracked neurons. We tested how well the classifier could decode the tone identity on any given trial from the same neurons on the same or different days. Tone identity was still successfully decoded with high accuracy from the subpopulation of tracked neurons on any given day (**Fig. 3f-h**), as was the case for the whole population of recorded cells (**Fig. 2e,f**). Tone identity could also be decoded well above chance by the same population on a different day, even with multiple days of separation (e.g., the classifier trained on P16 tracked cell activity could decode tones from P17-21 activity above chance, **Fig. 3g**). Although classifier decoding performance was unsurprisingly highest when trained and tested on tracked cell population activity from the same day, performance was still above the performance of classifiers trained on shuffled data across days, though with diminishing performance with larger train-test day offsets (**Fig. 3h**). This cross-day decoding performance from auditory cortical neuron activity in the first week of hearing is comparable to what has been described in the adult auditory cortex^50^ and to what we observed in our same animals recorded into adulthood. Altogether, these analyses of single-cell changes in tone-evoked activity in the first week of hearing demonstrate that despite some volatility and ‘drift’ at the level of individual neurons^50^, the postnatal auditory cortex possesses a stable and tonotopically organized representation of tone frequency from the earliest days of hearing.

### Emergence of USV responses in cells with high-frequency tuning

Having found that the postnatal auditory cortex reliably and stably represents the spectral features of simple tones from P14, finally we asked how the postnatal auditory cortex responds to and represents ultrasonic vocalizations (ultrasonic vocalizations, aka ‘calls’). Compared to pure tones, USVs are a more spectrotemporally complex, naturalistic, and behaviorally relevant class of acoustic stimuli, and their high ultrasonic spectral content often exceeds that of the range of pure tones that evoke activity in the auditory cortex of adult mice^38,40,51^. In contrast to the emergence of pure tone responses and other acoustic stimuli by P13-14, we observed a delayed onset of USV responses at P16-18 that peaked at the end of the third postnatal week in the same animals (**Fig. 4a-c**). In contrast to the developmental timecourse of auditory cortex responses to pure tones (**Fig. 2b**), the prevalence of USV-responsive cells starkly diminished over time, reaching adult-like levels by postnatal weeks 5-6 (**Fig. 4d**).

**Figure 4.**
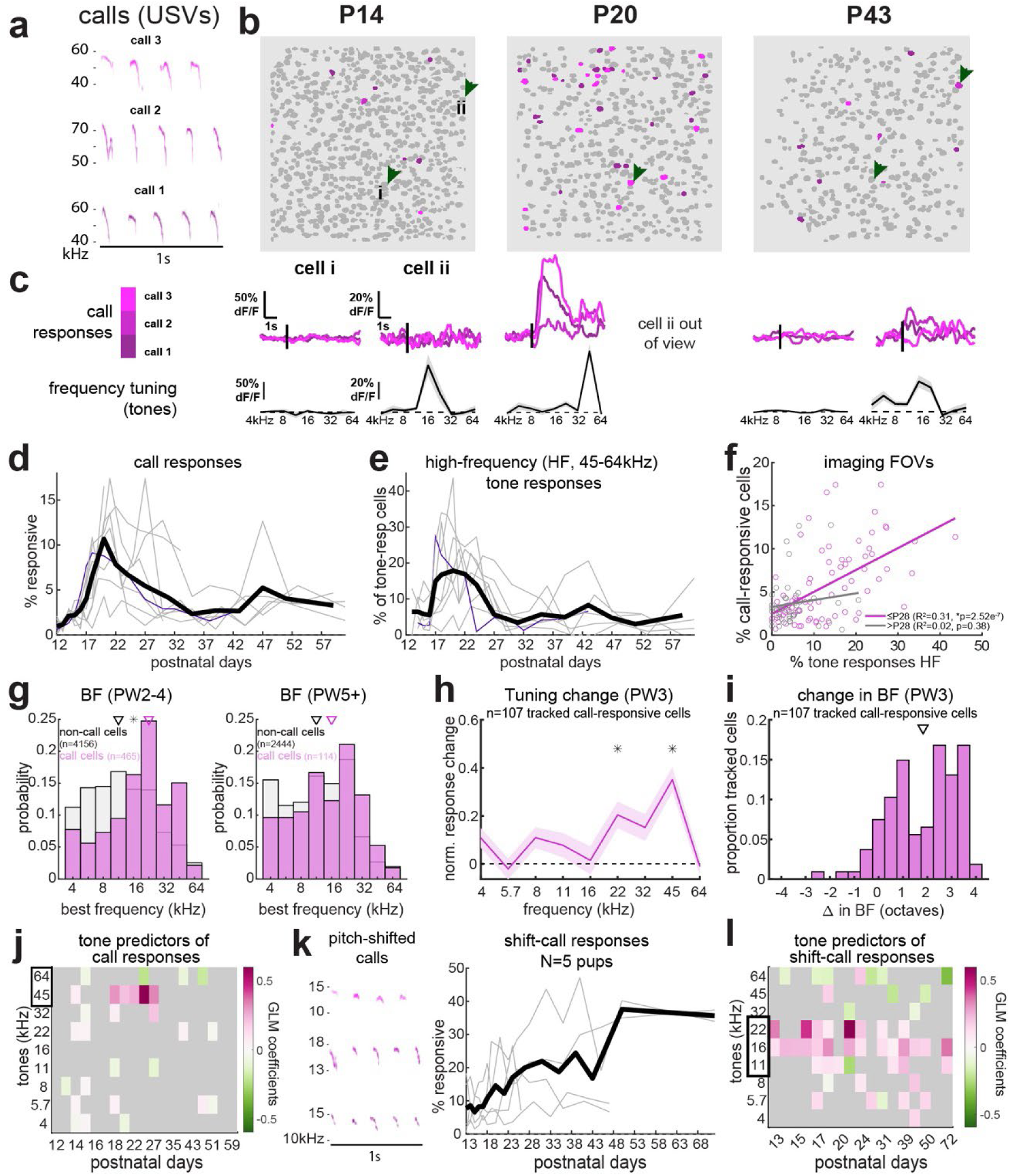
Call-responsive cells are tuned towards ultrasonic frequencies during early development, but not at later ages. **a**, Spectrograms of USVs used in experiments. **b**, Significant call-responsive cells across days in the same example animal as in prior figures (shade of pink corresponds to the call evoking the maximal median response). **c**, Top: call-evoked changes in fluorescence (dF/F) for two example cells. Bottom: mean tone-evoked dF/F response by tone frequency. **d**, Proportion of significant call-responsive ROIs across postnatal days for all animals (grey = individual animals; purple = example animal; black = mean across N=9 animals). **e**, Proportion of all tone-responsive cells with significant responses to high ultrasonic tones (45-64kHz). **f**, Correlations between cell-responsivity (as in **d**) and high-frequency tone responsivity (as in **e**) for all imaging FOVs for younger (<=P28) and older (>P28) ages. **g**, Comparison of distribution of best frequencies (BFs) between call-responsive and other tone-responsive cells at younger (left) and older (right) ages (p<0.001 and p=0.1144 respectively, Kolmogorov-Smirnov tests). **h,i**, Change in normalized tuning curves (**h**) and BF (**i**) between early-(P13-15) and late-(P17-21) postnatal week 3 for tracked, call-responsive cells. **j**, Significant tone-evoked response predictors of call-evoked responses in GLMs fit on all call- or tone-responsive neurons on each postnatal day. **k**, Left, spectrograms of pitch-shifted versions of the same three USVs as in **a** (down-shifted 2 octaves). Right, proportion of all cells responsive to these pitch-shifted calls in a separate group of animals (N=5). **l**, Significant coefficients of tone predictors for GLMs fit on responses to pitch-shifted calls.

We noticed that many of the emerging USV-responsive cells were also newly tuned to high-ultrasonic-frequency pure tones (cell i, **Fig. 4b**), specifically those in the same spectral range as the presented USVs (**Fig. 4a**). In fact, we noticed that the developmental time course of auditory cortex responses to these high-ultrasonic tones (45-64 kHz) was delayed relative to that of other tones and closely resembled the trajectory of USV responsivity (**Fig. 4e**). The unique developmental time course of auditory cortex responses to ultrasonic sounds was also evident in our linear classifier analyses, in which we observed peak classifier performance and discriminability of high-ultrasonic frequency tones P16-28 (**Fig. 2e,f**). Moreover, there was a high correlation between an imaging FOV’s proportion of call-responsive cells and its prevalence of high-ultrasonic tone responses (**Fig. 4f**), but only for FOVs recorded prior to postnatal week 5 (<=P28). Call-responsive cells at these later time points, while less prevalent, were still present and, consistent with prior descriptions^34–38,40^, were not especially tuned to ultrasonic frequency tones compared to the rest of the tone-responsive population (**Fig. 4g**; example cell ii in **Fig. 4b**). Not only did call-responsive cells at earlier ages exhibit spectral tuning for higher frequencies, but tracked call-responsive cells also demonstrated a shift towards higher frequency tuning (**Fig. 4h**) and best frequencies (**Fig. 4i**) over the third postnatal week, suggesting that cells may become responsive to USVs if and when their tuning curves expand and/or drift into the spectral range of USVs. To further test this, we fit a generalized linear model (GLM) to the call-evoked activity of all sound-(tone- or call-) responsive neurons, using responses to each of the nine presented pure tones as predictors. Consistent with our hypothesis, we found that neuronal responses to high-ultrasonic tones, especially 45 kHz, were highly predictive of call-evoked responses, but only during a limited developmental window (P18-28, **Fig. 4j**). This was still the case if we limited our GLM analysis to call-responsive neurons exclusively, or if we subsampled to equate the number of fitted cells’ responses over time. Therefore, the developmental changes in the predictive power of ultrasonic tone responses for call responses cannot be solely explained by developmental changes in overall call responsivity.

Non-spectral features of USVs, such as their temporal structure, are thought to be the more salient features of USVs for guiding recipient animals’ behavior^31,35^. Thus, in an additional set of animals (N=5), we sought to test whether the postnatal auditory cortex would process pitch-shifted vocalizations (spectral range down-shifted by two octaves) more like standard USVs given their identical temporal structure, or more like mid-range-frequency tones with similar spectral range. We found evidence for the latter; auditory cortical neurons showed higher, earlier, and more persistent responsivity to the pitch-shifted calls (**Fig. 4k**), and those responses were best predicted by tonal responses to mid-range frequencies across developmental time (**Fig. 4l**). Thus, at early ages, auditory cortical neurons’ responses to USVs are determined predominantly by their tonal receptive fields. However, around 4 weeks of age, the number of call-responsive neurons decrease at the same time that neurons’ spectral tuning and call responsivity become decoupled.

## Discussion

Here, we have demonstrated both the feasibility and the promise of utilizing two-photon calcium imaging to longitudinally monitor the functional maturation and plasticity of the postnatal auditory cortex in young mice. This approach has proven fruitful for measuring developmental changes in sensory representations of lower-level acoustic features (frequency of pure tones) as well as complex, ethological auditory objects (USVs) across the lifespan of individual animals, and how those representations at both population- and single-cell-levels change over days to weeks of postnatal development.

Our observations of population-level changes in response to pure tone stimuli are broadly consistent with past studies on auditory cortical activity in postnatal rodents^8,10,48,49^. The specific timing of the onset of tone-evoked activity we observe, as well as the expansion of the tonotopic map over the first few days of hearing, are likely driven by the cochlea^48^, which is itself in its final stages of maturation with decreasing sensitivity thresholds and increasing sensitivity range at the beginning of the third postnatal week^43,52,53^. Although we have more consistently observed tone-evoked responses at P13 in additional experiments for which we presented tones at higher intensities (85-90 dB SPL versus 70 dB SPL for all data presented herein), we still very rarely observed reliable sound-evoked activity at P12. Instead, at this time, most animals exhibited high degrees of spontaneous and highly correlated network activity, consistent with descriptions of pre-sensing cortex^46,54^. On one hand, a P13 onset of sound-evoked activity is somewhat later than expected based on prior descriptions of hearing onset occurring P10-P12^42,43^ and the thalamocortical auditory critical period starting at P12 in mice^55^. It is plausible that the invasive surgical procedure at such a young age (P10-11) slightly retarded auditory cortex and/or hearing maturation. Consistent with another recent study utilizing longitudinal imaging in young pups^14^, we observed a minor slowing in weight gain of pups undergoing surgery, which that other group found to correlate with functional development^14^. Nonetheless, the overarching similarities in the developmental time course of sound-evoked activity we observe with postnatal imaging of the auditory cortex to other studies employing different recording methodology would suggest that any developmental delays introduced by our surgical approach are minor, at most one day’s worth of postnatal development.

A major advantage of our longitudinal imaging approach is the ability to follow neurons across developmental time, which has allowed us to characterize how changes in sound representations at the level of individual neurons underlie large-scale maturation of the auditory cortex’s functional map. Our observations of many cells with changing tone responses (gains and losses of responses, as well as shifts in frequency tuning) at the same time that we could decode the frequency of presented tone from the same cells’ activity across multiple days above chance, are similar to what has been described in the adult auditory cortex^50^. Thus, it is notable that the developing auditory cortex exhibits nearly adult-like degrees of sensory map stability already in the first week of hearing. The fact that individual cells tend to either maintain their same best frequency or ‘drift’ to closely related ones likely reflects the stability of thalamocortical connectivity, which has been shown to be in established prior to hearing onset^42,55^. Thus, tuning drift is likely caused by synaptic plasticity of existing thalamocortical connections, which may change which tones elicit responses that are supra- or sub-threshold for detection with GCaMP8m^55,56^. In the future, this imaging approach might be applied to test whether biases in single-cell tuning drift underlies experience-dependent plasticity of the tonotopic map during the critical period^8,10,57^.

We also observed that responses in the auditory cortex to ultrasonic sounds – both ultrasonic tones as well as playbacks of ultrasonic vocalizations – are delayed and down-modulated with age relative to responses to other sounds. To our knowledge, this study is among the first to describe the developmental onset of auditory cortex responses to pup ultrasonic vocalizations (but see ref.^27^ for responses to adult vocalizations). At early ages, we found that vocalization responses were closely related to cells’ sensitivity to high-ultrasonic frequency tones coupling, which contrasts with ours and others’ observations of unrelated spectral and vocalization tuning at later ages^34–38^. These observations raise at least two key questions. First, what drives the unique developmental trajectory of auditory cortex responses to ultrasonic stimuli writ large? Second, how do sensory representations for ultrasonic tones and calls become decoupled with age? The delayed onset may be inherited from the cochlea, where there is further developmental maturation and remapping of the place code over the first few days of hearing such that hearing thresholds for high frequencies take longer to mature than for lower frequencies^53,58,59^. Place code remapping, whereby the basal region of the cochlea develops first but then becomes mechanically tuned to progressively higher frequencies^53,58,59^, could also account for the slight bias in tuning drift in auditory cortical neurons towards higher frequencies, especially among call-responsive neurons. On the other hand, a larger range of frequency representations has been described upstream in the auditory brainstem at earlier timepoints^60^, suggesting that the unique time course of ultrasonic representations in the auditory cortex may be a cortical phenomenon, perhaps due to maturing lateral inhibition^37,38,40,61^. This latter possibility could also address the question of how call responses and ultrasonic tuning become decoupled, as lateral inhibition has been suggested to modulate call responses in mouse dams^38,40^. Recording from upstream thalamocortical neurons and from cortical inhibitory neurons in response to ultrasonic stimuli will help to determine the drivers of developmental changes in cortical receptive field properties for ultrasonic tone and vocalization stimuli.

Altogether, our longitudinal imaging approach reveals the developmental progression of sound-evoked activity and sensory representations in the postnatal auditory cortex. By gaining long-term, optical access to the actively developing sensory cortex in an awake, sensing animal, this work paves the way towards deciphering the plasticity processes and circuit mechanisms that instruct our observed developmental changes in sensory representations.

## Methods

### Animals

Experiments were performed primarily with C57BL/6J mice (N=9 animals). A subset of additional experiments featured in **Fig. 4k,l** were performed on offspring of homozygous Ai9 (JAX 007909) females bred to homozygous VGAT-Cre (JAX 028862) males (N=5). All experiments were performed following NYU Grossman School of Medicine IACUC and NIH guidelines for proper animal care and use in research. One animal was excluded that showed poor sound responsivity and tonotopic organization throughout the course of imaging, likely because the imaged field of view (FOV) did not fall within the auditory cortex.

### Neonatal injections

Mouse pups were neonatally injected with an AAV to drive jGCaMP8m expression (AAV9-CaMKIIa-jGCaMP8m-WPRE in C57 mice, or AAV9-syn-jGCaMP8m-WPRE in VGAT-Cre; Ai9 mice) at P0-1. Neonate pups were cryoanesthetized on ice, secured in a stereotax (Kopf) with a custom neonate adapter, and the skin over the left auditory cortex was punctured with a glass micropipette lowered with a motorized manipulator (MP-285, Sutter Instruments). Approximately 200 nl of virus was pressure-injected at each of two depths (∼680μm and 280μm). Pups recovered on a heating pad and were returned to their home cages upon regaining color and movement.

### Cranial window and headpost implants

Cranial window surgeries were performed at P10-11. Pups were anesthetized with isoflurane (1-2% vaporized in oxygen at a rate of 2 liters/minute) and secured in a stereotax with a nose cone adapted for young mice (Model 934-B, Kopf). Anesthesia levels were closely monitored based on the animal’s breathing rate and response to toe pinch, and body temperature was maintained at 37° C on a heating pad. Briefly, the scalp was trimmed of hair, cleaned with betadine, and coated with lidocaine. After a patch of scalp was removed to expose the skull, which was then cleaned of connective tissue and scored so that a custom headpost could be secured on the skull over the right hemisphere with dental cement (Metabond). Upon drying, this headpost was then used to further secure the head in place for separation of the left temporalis muscle from the skull and for a small, 3 mm diameter craniotomy over the left auditory cortex, which was located based on skull and blood vessel landmarks. Because the skull is very thin at these young ages, this craniotomy was performed with a small-burr dental drill, applied extremely lightly and with saline-soaked sponges regularly applied to clean and wet the skull. Dexamethasone (2 mg/kg) was preemptively delivered subcutaneously at the start of the craniotomy. Upon creating a ∼3 mm ring of cracked skull, the skull was carefully pulled away with great care to leave the dura intact. A circular glass window (3 mm diameter, FST/Harvard apparatus, thickness 0.1 m) was situated over the exposed brain moistened with saline and pressed upon gently with a blunt needle while the edge of the window was sealed to the skull edge from the craniotomy with a mixture of cyanoacrylate glue and acrylic resin (Ortho-Jet powder). Edges of the skin were sealed to the remaining exposed skull with tissue adhesive (Vetbond, 3M), and more dental cement was used to cover the remaining exposed skull between the cranial window and headpost. Post-op analgesia included extended-release buprenorphine (Ethiqa-XR, 2.5 mg/kg and an additional dose of dexamethasone (2mg/kg), and pups recovered for ∼45 min on a heating pad before being returned to the home cage with their littermates and mother. Additional doses of dexamethasone were delivered for 2-4 days following surgeries. Because we found that maternal rejection was the primary barrier to recovery and survival in our pups, we routinely used the same dams that had demonstrated good care of implanted pups. We then co-housed new nulliparous females with those experienced dams and implanted pups prior to breeding them and using their pups for surgeries.

### *In vivo* two-photon calcium imaging

Imaging experiments commenced at P12, following full surgery recovery. Mice were stabilized inside a custom 3D-printed tube holder with a custom headpost holder (ThorLabs parts) under the two-photon microscope (Moveable Objective Microscope, Sutter Instruments outfitted with resonant-galvo scanners). Two-photon stimulation of GCaMP8m at 900 nm was achieved with a Mai Tai DeepSee HP ultrafast Ti:Sapphire laser (SpectraPhysics). We used ScanImage (MBF Bioscience) to collect images through a Nikon 20x objective over a 400-500μm field of view with 512x512 pixel resolution at a 30Hz frame rate. The same imaging field of view was located across days based on blood vessel landmarks, which then helped us to locate the same precise field of view by comparing to the mean two-photon images taken on prior days. We focused on superficial (layer 2/3) activity by focusing our imaging plane down to 160-200 μm beneath the cortical surface, although in two animals we recorded slightly deeper (250-300 μm). At the end of daily recordings, we also collected z-stacks (60-90 frames at each of 30-40 depths with 2 μm spacing).

Auditory stimulation was generated and delivered with a digital signal processor (RZ6, Tucker-Davis Technologies) driving an ultrasonic speaker (CD1, Tucker-Davis Technologies) positioned 10cm from the mouse’s right ear. Stimuli consisted of nine pure sine-wave tones (4-64 kHz, half-octave spacing, 70 dB SPL, 25 0ms duration including 10 ms rise-fall time) and playbacks of three ultrasonic vocalizations that have been used in other studies from our lab^34^. Tone stimulus intensity was calibrated with an ultrasonic microphone and ACOustical interface system (ACO Pacific) through the SigCalRP software (Tucker-Davis Technologies).

We typically imaged the same fields of view every day for at least one week (with rare exceptions), and then with increasing intermittence (e.g., every other day P18-22, every third day P24-30, etc.) for as long as windows remained clear and animals remained healthy. We also monitored animals’ weight to ensure that they were growing normally and conducted post-hoc histology to verify virus targeting to the auditory cortex and to confirm that there were no gross brain deformations caused by the cranial implants during development.

### Two-photon image processing

Two-photon imaging data collected as TIF files was preprocessed and motion-corrected using Suite2P^60^. ROIs corresponding to putative individual neurons were further curated manually with the Suite2P GUI. When available, we also registered our data to collected z-stacks in order to compute Z-plane position across frames. Frames in which Z-position deviated by more than 2 standard deviations from the mean position or by more than 10μm were excluded from subsequent analyses, as were full trials if over half of their baseline or stimulus periods consisted of bad motion frames. Average fluorescence from pixels corresponding to the same ROIs were extracted and neuropil-corrected (F_t_ = F_soma_-0.7*F_neuropil_), and converted to trial-wise dF/F traces. dF/F was calculated as (F_t_-F_0_)/F_0_ for each stimulus trial where F_0_ is the mean of the one-second period immediately prior to stimulus onset.

To identify ROIs that correspond to the same neurons across multiple days of imaging, we used Track2P^14^. While this algorithm compares consecutive pairs of days and excludes any ROIs that were not identified across both days, we modified this package in order to compare multiple day pairs with up to three imaging days of separation.

### Two-photon data analyses and statistics

Analyses and statistics were performed with custom MATLAB scripts. The criteria for being considered ‘significantly responsive’ to any given stimulus was a) dF/F exceeding 2 standard deviations above the mean of the pre-stimulus (1 second) period during the stimulus window on at least half of trials, and b) the mean dF/F over the stimulus window was significantly greater than the mean dF/F over the pre-stimulus period across trials (p<0.01, paired, one-tailed t-test). For each stimulus class (tones, calls), the ‘best stimulus’ (e.g., best frequency, BF) was defined as the stimulus that elicited the largest median dF/F response over the stimulus window and across trials. The response window for tones was 450ms long (given the 250 ms tone duration plus 200 ms to account for rise and decay times of GCaMP8m^41^). For calls – which consist of 3-4 syllables over the course of ∼1 second –, the response window was 1275 ms.

Between-animal comparisons over the course of development was made more challenging by the fact that not every animal was recorded for the same length of time or on the exact same postnatal days (especially at later timepoints when imaging became more intermittent). Thus, we grouped some postnatal days (particularly later days) together for between-animal comparisons. The edges (in postnatal days) of the time bins used for all analyses presented here were: [13 14 15 16 17 18 20 22 24 27 31 35 39 43 47 51 55 59 63].

To determine a given field of view’s ‘tonotopic axis’ (**Fig. 2**), we performed linear regression (MATLAB function fitlm) on cells’ BFs and their x-y coordinates and then computed the angle of that linear fit in the x-y plane. We then correlated cells’ position along that axis with their BF to obtain R^2^ metrics of tonotopy, i.e., the amount of variances in cells’ BF explained by their position along the tonotopic axis.

For tracked cell analyses over the course of the third postnatal week (**Fig. 3**), we binned imaging sessions over that period into two groups; ‘early’ (P13-15), and ‘late’ (P17-21). Tracked cells were classified as ‘lost response’, ‘gained response’, ‘persistent response’, or ‘no response’ based on whether they exhibited a significant response (see above) to any of the tonal stimuli on any day in each of these day ranges. Because cells were often significantly responsive on more than one imaging day within each group of days, for direct comparisons of ‘early’ versus ‘late’ tuning curves and BF we used the first ‘early’ day with a significant response and the last ‘late’ day. Tuning curves were normalized to the largest median response across tonal frequencies, and we computed the Pearson correlation coefficient (using MATLAB function corrcoef) between these normalized tuning curves among ‘persistent responsive’ cells to assess their degree of ‘stability’ versus ‘drift’.

A transition probability matrix was computed by counting the number of transitions from one BF (or lack of a significant tone-evoked response) to another for all cells that were tracked for at least two consecutive imaging days and exhibited significant tone responses at any point between P13-21. A shuffled distribution of transition probabilities was generated from 1000 iterations of shuffling the *n* x *m* matrix of BFs (0-9, 0=no response) with *n*=number of tracked cells and *m=*9 (9 days P13-21). Compared to this distribution, a real transition probability was deemed significant if the probability *p* of observing that same transition rate in the shuffled data was less than or equal to the critical value *a*, corrected with the Benjamini-Hochburg method to yield at most an expected 2.5% false discovery rate.

### Linear classifier analyses

We utilized linear discriminant analysis (MATLAB cvpartition and fitcdiscr functions) to train and test linear classifiers’ ability to decode the frequency of tone that elicited a particular pattern of activity across cells. For same-day classifiers in **Fig. 2**, we fit the models on the responses (median dF/F over stimulus window for each trial) of all recorded cells (with 100 repetitions of k-fold cross validation, k=10), whereas in **Fig. 3**, we used only those neurons that were tracked between that day and another day. For cross-day decoding analyses (**Fig. 3**), we identified all neurons that were tracked between a given pair of imaging days and used just those cells’ responses to train a classifier on one day and test decoding on the other day. Whenever there were unequal numbers of trials for a particular stimulus, we randomly subsampled from over-represented trial types to obtain equal trials numbers. We also trained classifiers on shuffled data in order to compare the decoding performance of classifiers trained on shuffled versus real data. ‘Classifier performance’ was computed as the proportion of all of the classifier’s guesses that were correct (i.e., predicted tone = actual tone). Statistically significant cross-day comparisons were determined with the Wilcoxen signed-rank test on shuffled versus real classifier performance across N=9 pups with the same Benjamini-Hochburg FDR correction method.

### GLM analyses

For each imaging day (or range of imaging days at later timepoints, as described above), a generalized linear model (GLM) was fit (using MATLAB fitglm function) on the median call-evoked responses of all significantly call- or tone-responsive cells, using their tone-evoked responses to each of the nine tonal frequencies as predictors. The coefficients of significant predictor variables are shown in **Fig. 4**. The same approach was taken to predict pitch-shifted (2 octaves down-shifted) call responses from tone responses among shifted call- and tone-responsive cells.

## Notes

### Competing Interest Statement

The authors have declared no competing interest.

### Funding

- National Institute on Deafness and Other Communication Disorders: DC012557
- Eunice Kennedy Shriver National Institute of Child Health and Human Development: HD088411
- National Institute of Neurological Disorders and Stroke: NS107616

